# Multi-location trials identify stable high yielding spring bread and durum wheat cultivars in Mexico

**DOI:** 10.1101/2022.10.17.512639

**Authors:** Jorge L Valenzuela-Antelo, Ignacio Benitez-Riquelme, Mateo Vargas-Hernandez, Julio Huerta-Espino, Alison R Bentley, Hector E Villaseñor-Mir, Francisco J Pinera-Chavez

## Abstract

Determining the stability and consistency of grain yield performance requires accurate evaluation of genotypes in different environments. In Mexico, annual national spring wheat irrigated trials are conducted to assess elite bread and durum wheat performance in different testing environments (TEs) in the main wheat-growing areas. These trials provide data supporting release of new cultivars and aim to also address Mexican wheat value chain grain needs. In this study we analyzed 30 bread and durum wheat trial results from the 2012/13 and 2013/14 growing cycles conducted across TEs in northwest, north and central Mexico. Environmental variability (location, sowing timing, and irrigation schemes) across the national spring wheat irrigated trials enabled genotype by environment interaction to be effectively evaluated. We identified genotypes with high and stable grain yield across TEs of the wheat-growing areas of Mexico. The bread cultivars Bacorehuis F2015 and Borlaug100 F2014, and the durum cultivars Barobampo C2015, CONASIST C2015 and Anatoly C2011 were high yielding and gave stable performance in most of the TEs. This analysis demonstrates the utility of multi-year, multi-environment testing and analysis to identify improved wheat cultivars to meet wheat production demand in Mexico.

## INTRODUCTION

Wheat breeding aims to develop material with high and stable yield potential, resilience to biotic and abiotic stress and superior end-use quality (Crespo-Herrera et al., 2017). Selection of cultivars with these attributes requires multi-environment testing under near optimum conditions in order to select for yield potential as well as specific screening for abiotic and biotic stresses in each environment (Braun et al., 1996). The resulting cultivars should possess yield potential along with favorable gene accumulation for broad spectrum resistance to diseases and tolerance to abiotic stresses. Selection for target population of environments (TPEs) is key to designing strategies for crop improvement (Atlin et al., 2000). A recen analysis of spring wheat performance in india defined TPEs, showing that there is opportunity to further refine breeding targets based on current and future environmental parameters (Crespo-Herrera et al., 2021).

Identifying outstanding genotypes is complicated by the genotype by environment interaction (GEI) because ranking of the best genotypes can vary between environments (Asfaw *et al*., 2009). Strategies such as stratification of environments into mega-environments with specific biotic and abiotic stress profiles can be applied to reduce this difficulty (Yan et al., 2000). Several statistical methodologies have also been developed to identify superior high yielding and stable cultivars, termed “reliable genotypes” (Annicchiarico, 2002).

Static stability (type 1 or biological) refers to a constant expression of a character in relation to the environmental variation (phenotypic variation across environments = 0). Dynamic stability (type 2 or agronomical) refers to a character expression pattern which is parallel to the environmental variation, indicating the absence of GEI (Becker and Leon, 1988; Annicchiarico, 2002). Static stability is useful with characters (quality, disease resistance, tolerance to abiotic stresses, etc.) that have to be permanently fixed across environments (Becker and Leon, 1988). However, it is also commonly associated with poor response capacity to environmental changes and targeting static stability alone can lead to low yield potential. Therefore, dynamic stability is more appropriate to assess grain yield stability (Lin *et al*., 1986) as it allows for selection of environmental responsiveness.

One of the most commonly used methodologies for evaluating dynamic stability is the linear regression coefficient (*b_i_*) from the environmental mean for a given trait (Eberhart and Russell, 1966). Based on this methodology, a stable genotype would have a *b_i_* = 1, whereas genotypes that have *b_i_* ≠ 1 are sensitive to environmental changes. A genotype with *b_i_* > 1 tends to respond to high-yielding environments, whereas *b_i_* < 1, usually perform better in low-yielding environments (Eberhart and Russell, 1966). Multivariate methods, namely GGE biplots have arisen as an alternative to evaluate the GEI in multi-environmental trials (Kroonenberg, 1995; Tinker et al., 2006). This analysis uses the genotype (G) and genotype by environment (GE) effects to create graphical representations of genotype performance across multiple environments (Yan et al., 2000). GGE biplots are constructed using the first two principal components (PC1 and PC2) obtained from a single value decomposition of environment-centered yield data (Yan et *al*., 2000). The primary use of GGE biplots is to identify cultivars that have the best performance across a set of environments (Yan *e*t al., 2000).

In wheat, most previous studies have aimed to identify similarities and correlations between yield and yield components (Cossani, et al., 2011; Marti and Slafer, 2014). In this study we aimed to assess the contrasting yield performance and stability of performance of elite bread and durum wheat under irrigated conditions in Mexico. This builds on previous work to assess yield stability under rainfed conditions in central Mexico (Rodríguez-Pérez et al., 2002). The specific objectives were to: a) analyze the effect of environmental variation on the performance of genotypes from national spring bread and durum wheat in differente testing envrionments (TEs); b) compare phenotypic stability between bread and durum wheat genotypes; and c) identify high yielding stable cultivars.

## MATERIALS AND METHODS

### Trials and germplasm

Genotypes evaluated in the national spring wheat irrigated trials (denoted ENTRI) of the National Wheat Program of the National Forestry, Agricultural and Livestock Research Institute (INIFAP) were grown in the 2012/13 winter growing season in 13 locations and in 2013/14 in 16 locations with different testing envrionments (TEs) in northwest, north and central Mexico. The genotypes tested included eight bread and eight durum wheat cultivars (Table 1) which were planted in eight representative locations from the three major wheat-growing regions in Mexico (Table 2). Four TEs were defined using a combination of sowing dates (optimum and late) and irrigation management (full and reduced), resulting in a total of 30 trials (Table 2). The trials were determined by the following factors: growing season, location, irrigation management and in some cases sowing date. All experiments were sown using a randomized complete block design with two replicates. Experimental units consisted of plots of 3 m length and 1.2 m width with four plant rows. Agronomic, pest, and disease control management followed standard INIFAP procedures for each wheat-growing region. Grain yield (at 12% humidity) was obtained by harvesting the entire experimental plot after maturity.

**Table 1.** Cultivars included in the 2012-2013 and 2013-2014 trials.

| Cultivar name | Spring wheat type |
| --- | --- |
| Tacupeto F2001 | Bread |
| Kronstad F2004 | Bread |
| Urbina S2007 | Bread |
| Roelfs F2007 | Bread |
| Villa Juarez F2009 | Bread |
| Borlaug 100 F2014 | Bread |
| Conatrigo F2015 | Bread |
| Bacorehuis F2015 | Bread |
| Gema C2004 | Durum |
| CIRNO C2008 | Durum |
| Sawali Oro C2008 | Durum |
| Cevy Oro C2008 | Durum |
| Anatoly C2011 | Durum |
| Movas C2009 | Durum |
| CONASIST C2015 | Durum |
| Barobampo C2015 | Durum |

**Table 2.** Summary of the trials established in the 2012/13 and 2013/14 growing seasons in Mexico, including four testing envrionments (TE). Full irrigation with optimum sowing date (1), reduced irrigation with optimum sowing date (2), full irrigation with late sowing date (3) and reduced irrigation with late sowing date (4).

| Code | Growing cycle | Location | Longitude and Latitude | Region | TE<sup>ab</sup> |
| --- | --- | --- | --- | --- | --- |
| E1 | 2012-13 | Cd. Obregon, Sonora | 27°22' N, 109°55' W | Northwest | 1 |
| E2 | 2012-13 | Cd. Obregon, Sonora | 27°22' N, 109°55' W | Northwest | 2 |
| E3 | 2012-13 | Cd. Obregon, Sonora | 27°22' N, 109°55' W | Northwest | 3 |
| E4 | 2012-13 | Cd. Obregon, Sonora | 27°22' N, 109°55' W | Northwest | 4 |
| E5 | 2012-13 | Mexicali, Baja California | 32°32' N, 115°24' W | Northwest | 1 |
| E6 | 2012-13 | Los Mochis, Sinaloa | 25°45' N, 108°48' W | Northwest | 1 |
| E7 | 2012-13 | Los Mochis, Sinaloa | 25°45' N, 108°48' W | Northwest | 2 |
| E8 | 2012-13 | Roque, Guanajuato | 20°34' N, 100°48' W | Central | 1 |
| E9 | 2012-13 | Roque, Guanajuato | 20°34' N, 100°48' W | Central | 2 |
| E10 | 2012-13 | Roque, Guanajuato | 20°34' N, 100°48' W | Central | 3 |
| E11 | 2012-13 | Roque, Guanajuato | 20°34' N, 100°48' W | Central | 4 |
| E12 | 2012-13 | La Barca, Jalisco | 20°52' N, 102°42' W | Central | 1 |
| E13 | 2012-13 | Delicias, Chihuahua | 28°10' N, 105°29' W | North | 1 |
| E14 | 2012-13 | Rio Bravo, Tamaulipas | 25°57' N, 98°00' W | North | 1 |
| E15 | 2012-13 | Zaragoza, Coahuila | 28°35' N, 100°54' W | North | 1 |
| E16 | 2013-14 | Cd. Obregon, Sonora | 27°22' N, 109°55' W | Northwest | 1 |
| E17 | 2013-14 | Cd. Obregon, Sonora | 27°22' N, 109°55' W | Northwest | 2 |
| E18 | 2013-14 | Cd. Obregon, Sonora | 27°22' N, 109°55' W | Northwest | 3 |
| E19 | 2013-14 | Cd. Obregon, Sonora | 27°22' N, 109°55' W | Northwest | 4 |
| E20 | 2013-14 | Mexicali, Baja California | 32°32' N, 115°24' W | Northwest | 1 |
| E21 | 2013-14 | Los Mochis, Sinaloa | 25°45' N, 108°48' W | Northwest | 1 |
| E22 | 2013-14 | Los Mochis, Sinaloa | 25°45' N, 108°48' W | Northwest | 2 |
| E23 | 2013-14 | Roque, Guanajuato | 20°34' N, 100°48' W | Central | 1 |
| E24 | 2013-14 | Roque, Guanajuato | 20°34' N, 100°48' W | Central | 2 |
| E25 | 2013-14 | Roque, Guanajuato | 20°34' N, 100°48' W | Central | 3 |
| E26 | 2013-14 | Roque, Guanajuato | 20°34' N, 100°48' W | Central | 4 |
| E27 | 2013-14 | La Barca, Jalisco | 20°52' N, 102°42' W | Central | 1 |
| E28 | 2013-14 | Rio Bravo, Tamaulipas | 25°57' N, 98°00' W | North | 1 |
| E29 | 2013-14 | Zaragoza, Coahuila | 28°35' N, 100°54' W | North | 1 |
| E30 | 2012-13 | La Barca, Jalisco | 20°52' N, 102°42' W | Central | 2 |

<sup>a</sup>Full irrigation, four irrigation events after sowing; reduced irrigation, 3 irrigation events between sowing and flowering.
<sup>b</sup>Optimum sowing date was performed within the recommended sowing dates for each location, and the late sowing date was performed 15 days after the recommended sowing date.

### Statistical analysis

The Restricted Maximum Likelihood procedure was implemented using the META-R program to calculate variance components and broad-sense heritability estimates for combined analysis and individual environments (trials) considering all factors as random effects (Alvarado et al., 2020) (Eq. 1).

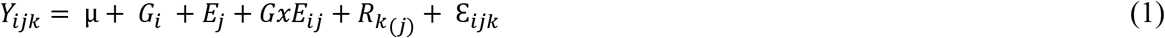

Where *y_ijk_* is the trait of interest; μ the mean effect; *G_i_* is the genetic effect of the *i^th^* genotype; *E_j_* is the effects of the *j^th^* environment; *GxE_ij_* is the genotype-by-environment interaction effect; *R_k_* is the effect of the *k^th^* replicate nested within the *j^th^* environment; and 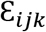 is the residual.

Least significant difference (LSD _0.05_) was used to compare the Best Linear Unbiased Predictor (BLUP) in the genotype combined analysis (Eq. 2).

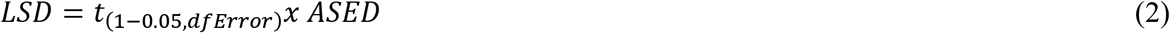

Where the *t* is the cumulative Students’s distribution with a level of significance of 5 %, *dfError* is the degrees of freedom of the error, and *ASED* is the average standard error of the differences in means (Alvarado et al., 2020).

Heritability estimates for individual environment and combined analysis were calculated as in equation 3 and 4, respectively.

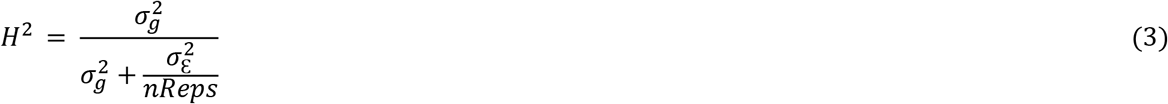

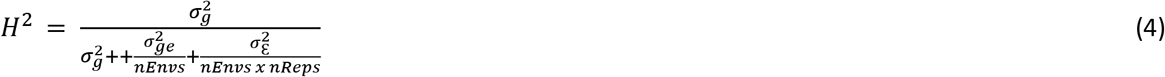

Where *H*^2^ is broad-sense heritability, 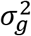, 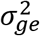 and 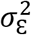 are genotype, genotype by environment and error variances, respectively, nReps is the number of replications, and *nEnvs* is the number of environments. Environments with *H^2^* below 0.05 were eliminated from the analysis (resulting in the removal of trial E30 (Table 2)).

The Eberhart and Russell (1966) stability parameter bi, and the GGE biplot analysis were used to determinte staibility and GEI. The GGE biplots were generated using the first two principal components, which explained most of the G and GE variability based on the model (Eq. 5) (Yan & Tinker, 2005):

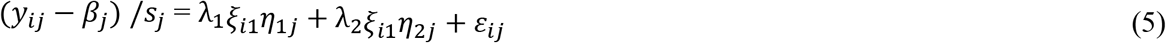

where *y_ij_* is the mean of the *i* genotype in the *j* environment; *β_j_* is the mean yield in environment *j*; *s_j_* is the standard deviation in the environment *j*; *λ*_1_ and *λ*_2_ are the singular values of PC1 and PC2, respectively; *ξ*_*i*1_ and *ξ*_*i*2_ are the eigenvectors of the genotype *i* for PC1 and PC2, respectively; and *η*_1*j*_ and *η*_2*j*_ are the eigenvectors of environment *j* for PC1 and PC2, respectively; and ε_jj_ is the residual of the genotype i and environment j. Equation 5, is reorganized to obtain the principal component scores of *i*^th^ genotype and *j*^th^ environment:

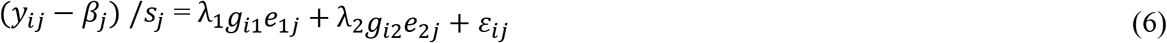

Where *l*= 1, 2; *g_il_* is the PC scores for genotype *i*^th^ (*ξ_il_*), and *e_lj_* PC scores for the *j*^th^ environment (*η_lj_*).

The Eberhart and Russell (1966) analysis of variance (ANOVA) and stability parameters were obtained using the “plantbreeding” package (Umesh, 2014) implemented in R (R Core Team, 2020). The GGE biplots “which-won-where” and “means vs. stability” were created using the “gge” and “GGEBiplots” packages in R (Dumble et al., 2017; Wright and Laffont, 2018; R Core Team, 2020). The genetic correlation among environments using the dendrogram and biplot were performed using META-R (Alvarado et al., 2020). The biplots were generated using the 16 genotypes in all 29 trials to determine major testing environments based on the grain yield genotypic performance and to identify genotypes with high and stable yield.

## RESULTS

### Environmental and GEI variation impacts yield performance

A large portion of grain yield variation was explained by the environment (trials), followed by GEI and genotype. The combined analysis of variance showed statistically significant differences *(P* < 0.01) for all sources of variation (Supplementary Table S1). The per environment grain yield varied significantly, ranging from 8.3 t ha^-1^ in E5 (TE 1: Mexicali full irrigation with optimum sowing date in 2012/13) to 1.52 t ha^-1^ in E11 (TE 4: Roque reduced irrigation and late planting in 2012/13) (Table 3). Significant differences were also found in the variance of individual environments for nine out of the 29 environments (ranging from *P* < 0.01 and < 0.001). Heritability estimates for each experiment ranged from 0.09 – 0.93 but 20 environments had estimates > 0.5, the coefficient of variation (*CV*) ranged from 3.10 – 23.2 % with 22 environments having a *CV* < 15 %. Most of the high yielding environments were located in the northwestern region of Mexico with lower yielding environments in the Central region (Table 3).

**Table 3.** Variation for individual environment (as per Table 2) variance components, broad-sense heritability, grain yield means and coefficient of variation in 29 environments.

| Trial | Variance components |  | Heritability | Grain yield mean (t ha<sup>-1</sup>) | Coefficient of variation (%) |
| --- | --- | --- | --- | --- | --- |
|  | Genotype | Residual |  |  |  |
| E1 | 0.239* | 0.178 | 0.72 | 7.95 | 5.30 |
| E2 | 0.149* | 0.158 | 0.65 | 5.58 | 7.10 |
| E3 | 0.218*** | 0.044 | 0.90 | 6.69 | 3.10 |
| E4 | 0.560 | 0.110 | 0.48 | 4.67 | 7.35 |
| E5 | 0.490 | 0.820 | 0.54 | 8.30 | 10.92 |
| E6 | 0.850 | 0.140 | 0.54 | 5.82 | 6.51 |
| E7 | 0.080 | 0.140 | 0.54 | 5.82 | 6.51 |
| E8 | 0.370* | 0.340 | 0.69 | 3.95 | 14.83 |
| E9 | 0.190 | 0.330 | 0.54 | 2.64 | 21.61 |
| E10 | 0.230** | 0.130 | 0.79 | 3.65 | 9.77 |
| E11 | 0.370 | 0.050 | 0.93 | 2.12 | 10.99 |
| E12 | 0.030 | 0.240 | 0.22 | 2.52 | 19.30 |
| E13 | 0.510 | 1.620 | 0.39 | 5.48 | 23.21 |
| E14 | 0.180 | 0.470 | 0.44 | 3.28 | 21.01 |
| E15 | 0.480 | 0.690 | 0.58 | 3.59 | 23.15 |
| E16 | 0.030 | 0.460 | 0.12 | 5.20 | 13.09 |
| E17 | 0.030 | 0.190 | 0.27 | 4.59 | 9.48 |
| E18 | 0.040 | 0.110 | 0.44 | 4.53 | 7.39 |
| E19 | 0.150* | 0.100 | 0.75 | 5.06 | 6.37 |
| E20 | 0.270 | 1.310 | 0.30 | 7.74 | 14.76 |
| E21 | 0.300* | 0.220 | 0.74 | 6.13 | 7.63 |
| E22 | 0.480 | 0.570 | 0.63 | 5.45 | 13.86 |
| E23 | 0.910 | 0.230 | 0.89 | 5.94 | 8.01 |
| E24 | 0.020 | 0.550 | 0.09 | 4.82 | 15.38 |
| E25 | 0.130 | 0.270 | 0.50 | 4.58 | 11.31 |
| E26 | 0.140* | 0.130 | 0.69 | 3.94 | 9.16 |
| E27 | 0.460 | 0.680 | 0.58 | 5.78 | 14.26 |
| E28 | 0.250 | 0.290 | 0.63 | 3.52 | 15.35 |
| E29 | 0.400* | 0.41 | 0.66 | 4.43 | 14.50 |

\*, \*\*\*, Statistically significant with $P < 0.05$ and $< 0.001$ , respectively.

The genetic correlations between trials were also determined (Figure 1) showing that trials that belong to the northern region clustered together (Figure 1a in blue) and central region trials located mostly in one of the two mega-clusters (except for E27) (Figure 1a in red). Interestingly, for the northwestern region no consistent pattern was observed (Figure 1a in yellow).

**Figure 1.**
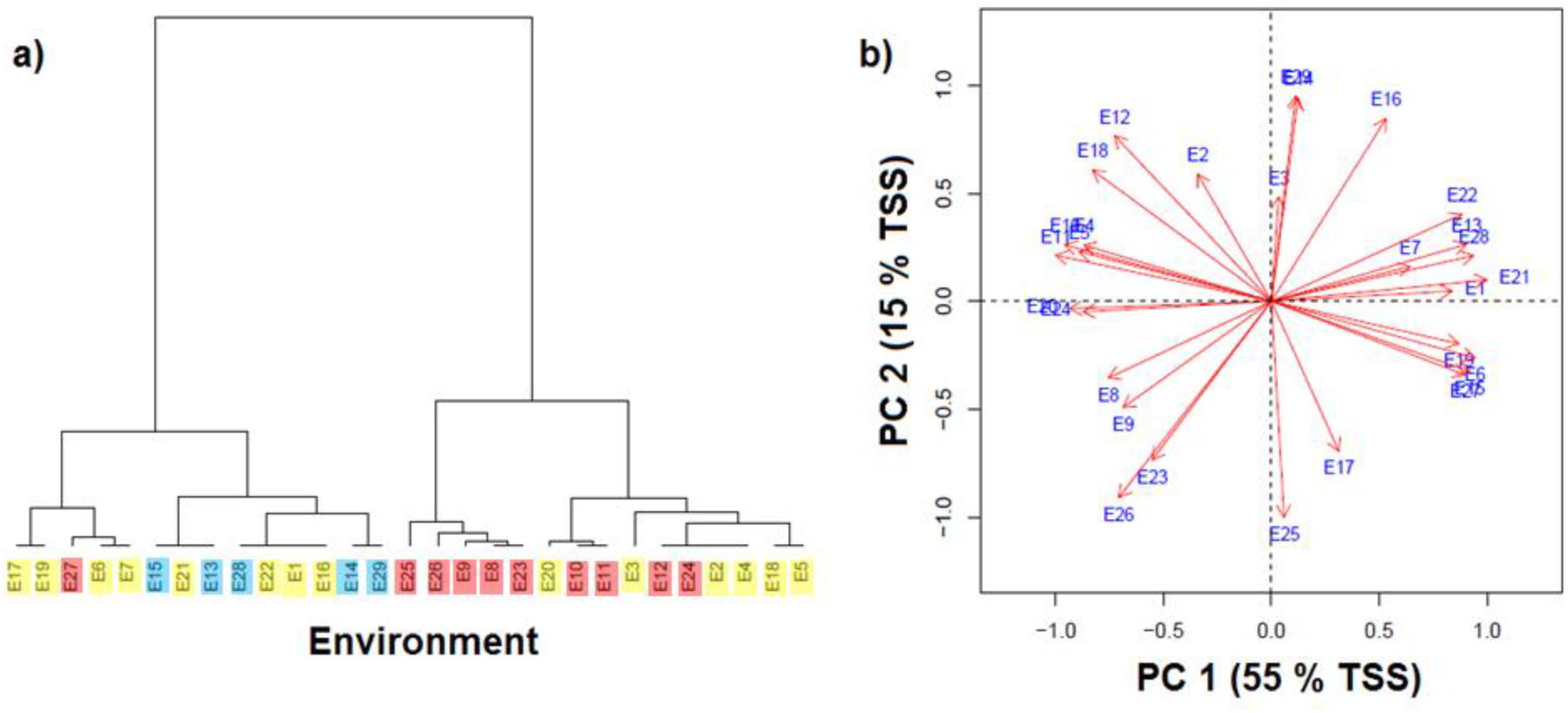
Clustering of trials: (a) dendrogram (northwest in yellow, north in blue and central in red)) and (b) biplot using the two principal components for environments clustering according their genetic correlation. See Table 2 for trial code.

### Stability assessment identifies genotypes with high potential across environments

Genotype rankings differed across environments, justifying the evaluation of cross-environment yield stability. The Best Linear Unbiased Predictors (BLUPs) per genotype (Table 4) demonstrate overall variation in grain yield differences. The bread wheat cultivars Bacorehuis F2015 and Borlaug100 F2014 and the durum wheat cultivars Barobampo C2015, Conasist C2015, and Anatoly C2011 had an overall grain yield of > 5.1 t ha^-**1**^.

**Table 4.** Grain yield overall Best Linear Unbiased Predictor (BLUP) and stability parameters *(b_i_)* (Eberhart and Russell, 1966) for bread and durum wheat cutlivars.

| Cultivar | BLUP (t ha<sup>-1</sup>) | Stability parameters ($b_i$) |
| --- | --- | --- |
| **Bread wheat** |  |  |
| Bacorehuis F2015 | 5.30<sup>a</sup> | 1.00† |
| Borlaug100 F2014 | 5.14<sup>ab</sup> | 0.87† |
| Conatrigo F2015 | 4.98<sup>bcd</sup> | 1.01† |
| Villa Juarez F2009 | 4.95<sup>bcd</sup> | 1.05§ |
| Roelfs F2007 | 4.91<sup>cd</sup> | 1.02† |
| Kronstad F2004 | 4.85<sup>de</sup> | 0.92† |
| Tacupeto F2001 | 4.66<sup>ef</sup> | 1.10† |
| Urbina S2007 | 4.57<sup>f</sup> | 0.84‡ |
| **Durum wheat** |  |  |
| Barobampo C2015 | 5.31<sup>a</sup> | 1.10† |
| CONASIST C2015 | 5.14<sup>ab</sup> | 0.91† |
| Anatoly C2011 | 5.13<sup>abc</sup> | 1.04† |
| CIRNO C2008 | 4.97<sup>bcd</sup> | 1.12† |
| Gema C2004 | 4.96<sup>bcd</sup> | 0.89† |
| Cevy Oro C2008 | 4.90<sup>d</sup> | 0.93† |
| Sawali Oro C2008 | 4.90<sup>d</sup> | 1.22† |
| Movas C2009 | 4.76<sup>def</sup> | 0.94† |
| **Overall LSD<sub>(0.05)</sub>** | 0.226 |  |

LSD, least significant difference ( $P = 0.05$ ) for all the cultivars; different letters indicate statistical differences; †, indicates stable genotypes ( $b_i = 1$ ); § indicates genotypes that perform better in favorable environments ( $b_i > 1$ ), ‡ indicates genotypes that perform better in unfavorable environments ( $b_i < 1$ ).

The stability analysis ANOVA (Eberhart and Russell, 1966) indicated that genotypes and GE (linear) are statistically different, meaning that there are differences among the cultivars regression from the environmental index (Supplementary Table S2). For the bread cultivars, only Villa Juarez F2009 and Urbina S2007 showed regressions coefficients different from 1 (Table 4, see Supplementary Table S2 for singifcant diferences). Therefore the latest released genotypes, Bacorehuis F2015, Barobampo C2015, CONASIST C2015, Borlaug100 F2014, along with Anatoly C2011 (Table 4) met the previously defined requirements of a reliable genotype (Annicchiarico, 2002). All the durum cultivars showed regressions coefficients from the environmental index statistically similar to *b_i_* =1 (Table 4).

The GGE analyses allowed visualization of the best genotype for each environment (Yang et al., 2009) (Figure 2) as well as for identification of the most stable genotype (Yan, 2001) (Figure 3). The percentage of the total variation explained by PC 1 and PC 2 of the GGE model was 54% and the bipot was divided into six sectors delimited by gray dotted lines in Figure 2 (S1 – S6). The 29 environments were grouped in sectors S1, S4, S5 and S6, vectors for northwest and central regions were located in sector S5 and north region vector was located in sector S6. TE 1 (full irrigation with optimum sowing date) was distributed in sectors S1, S4, S5 and S6, TE 2 (reduced irrigation with optimum sowing date) in sectors S5 and S6, TE 3 (optimum irrigation with late sowing date) in sectros S4 and S5 and TE 4 (reduced irrigation with late sowing date) in sectors S4, S5 and S6.

**Figure 2.**
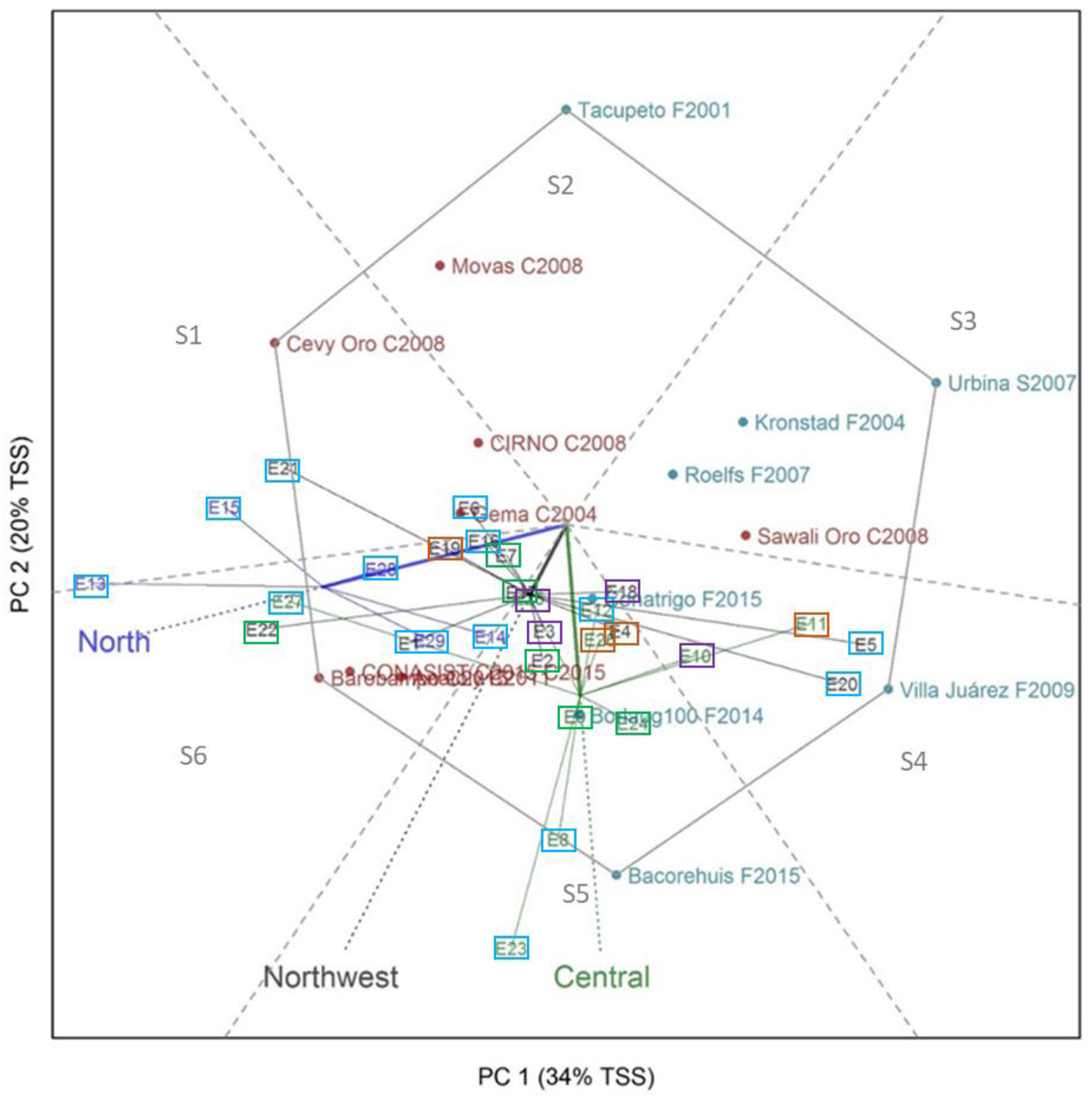
Which-Won-Where biplot with a polygon view obtained from the first two principal components (PC 1 and PC 2) for grain yield of bread (blue) and durum (red) wheat established during winter-spring of 2012/13 and 2013/14 at north (blue navy), northwest (black) and central (green) Mexico. Blue, green, purple and red squares indicated TEs 1, 2, 3 and 4, respectively. See Table 2 for trial code and TEs.

**Figure 3.**
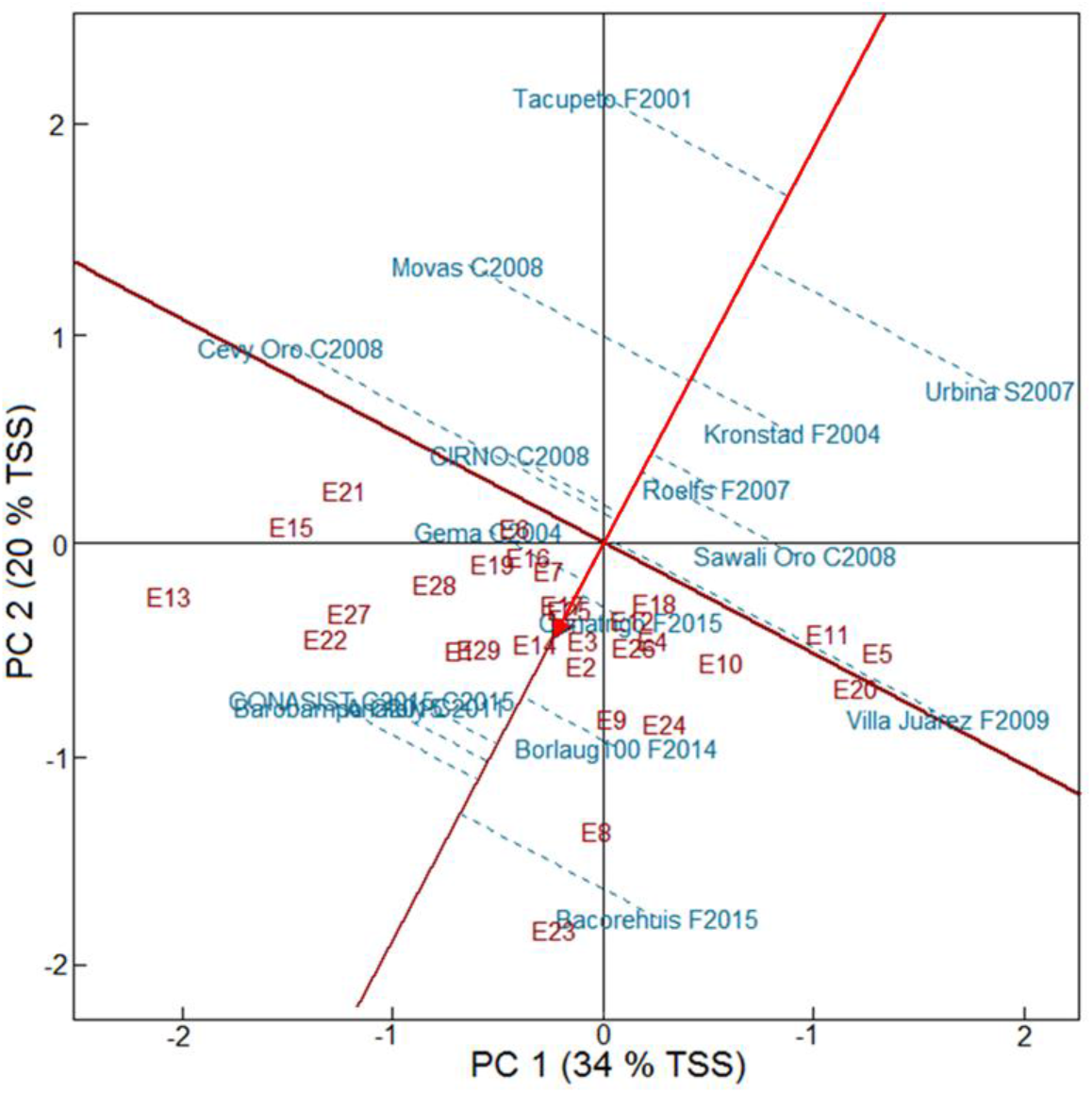
GGE biplot for stability versus grain yield obtained from the first two principal components (PC 1 and PC 2) for grain yield of bread and durum wheat established during winter-spring of 2012/13 and 2013/14 at the north, northwest and central Mexico. See Table 2 for trial code.

The GGE biplot in Figure 2 indicated highest yielding genotypes located at the polygon vertices for each sector. Bacorehuis F2015 was the highest yielding cultivar for the northwest and central regions (sector S5), whereas Barobampo C2015 was the highest yielding cultivar in the northern region (sector S6). Villa Juarez F2009 was the highest yielding cultivar in the sector S4 and Cevy Oro C2008 was the highest yielding in sector S1. Cultivars in vertices of sectors S2 and S3 had poor performances in most environments.

Another feature of the GGE analysis is the comparison of means vs stability (Figure 3) in which the red arrowed line indicates the average environment coordination (AEC) abscissa that points to the higher mean genotype. The dashed blue lines are the projection from the AEC coordinate, for which shorter lines mean higher grain yield stability (Yan et al., 2007). The results obtained corroborate the findings of the Eberhart and Russell (1966) stability analysis (Table 4) indicating that Conatrigo F2015, Roelfs F2007 and Gema C2004 are among the most stable cultivars given the short projection from the AEC ordinate. Barobampo C2015,

CONASIST C2015, Anatoly C2011, Borlaug100 F2014, CIRNO C2008, Sawali Oro C2008, and Kronstad F2004 also had relatively short projection and therefore classified as stable. In Figure 3, Bacorehuis F2015 was also identified as the high yielding cultivar but was unstable. When comparing the number of stable genotypes, a greater number of durum wheat cultivars were stable compared to bread wheat with a frequency of six and four, respectively.

## DISCUSSION

Previous work has reported that bread wheat yields more in high-yielding potential conditions whilst durum wheat is more productive in marginal conditions (Marti and Slafer, 2014). However, further reports have shown that durum cultivars have superior performance in high yielding conditions (full irrigation and/or optimal sowing date) and are similar to bread cultivars under marginal conditions (reduced irrigation and/or late sowing date) (Valenzuela-Antelo et al., 2018).

When evaluating yield performance and stability, we found differential expression of yield in different testing environments in Mexico. In addition, the phenotypic plasticity, or magnitude of response was different depending on the genotype. Studies have shown that higher yield stability is due to greater responsiveness, attributed to the ability of some genotypes to modulate yield components (Cossani et al., 2011; Sadras and Slafer, 2011). In both bread and durum wheat, the two major yield components that affect yield stability are the number of grain per m^2^ and grain size (Marti and Slafer, 2014). These authors showed that bread wheat tends to have higher grain per m^2^, whereas durum tends to has bigger grain. Lopes et al. (2012), found that the genetic gain in bread wheat elite lines from the International Maize and Wheat Improvement Center (CIMMYT) was linearly associated with increments in grain size. This linear association between high grain yield and grain size could be one of the reasons explaining similar stability for grain yield reported in the present study, as the tested cultivars are the product of a INIFAP-CIMMYT breeding network (Villaseñor-Mir, 2015).

Given the similar yield responsiveness seen in this study, differences in yield components between bread and durum wheat in Mexico are lower than previously reported in other regions (Pfeiffer et al., 2001). In addition, similar grain yield stability could be the result of substantial breeding efforts made to improve spike fertility, which is a determinant factor that affects yield when genotypes are subject to different photoperiod and temperature regimes (Reynolds et al., 2012). Further work using defined genetic stocks and/or material developed for enhanced physiological parameters is required to dissect the drivers of the yield stability identified in this study.

In this study we used two methods to determine stability: Eberhart and Russell (1966) stability parameters and GGE biplots. Both identified a similar set of stable cultivars, as has been previously reported (Alwala et al., 2010; Blanche et al., 2007). We also found concordance in the in the sector groups in the “Which-Won-Where biplot” and the dendrogram genetic correlations between trials. This highlights three points: firstly, the trials simulating testing environments (TEs) with a combination of irrigation level and sowing date (E1 – E7 and E16 – E22) in the three northwest region locations (Obregon, Mexicali and Los Mochis) were highly diverse according to the GEI of the genotypes and correlated with other trials in other regions. Secondly, the testing environments from the north and central regions tended to cluster together. Thirdly, the “Which-Won-Where biplot” showed a clear trend in the adaptation by wheat type with the durum cultivars tending to perform better in the northern environments, whereas the bread wheat cultivars performed well in the central region environments. Durum cultivars have dominated northwest wheat growing areas due to their great yield potential, resitance to karnal bunt and rapid adoption of superior varieties by farmers (Fischer et al., 2022). These findings indicate that a relative low number of trials targeting the TEs used in this study in the northwestern region of Mexico are sufficient for the identification of high-yielding and stable cultivars. In contrast, TEs trials in the northern and central regions may maximize the identification of superior cultivars with local rather than broad adaptation.

The cultivars tested and analyzed in this study had outstanding performance across TEs and were the best performers in the national spring wheat irrigated trials (ENTRI) prior to their release (Villaseñor-Mir, 2015). This indicates that in most cases they have accumulated favorable alleles that permit them to respond consistently across environments. The analysis methods used in this study were useful to identify outstanding genotypes with high yield potential and stability to provide consistent productivity in Mexican wheat production environments.

Overall, the bread cultivars Bacorehuis F2015 and Borlaug100 F2014, and the durum cultivars Barobampo C2015, CONASIST C2015 and Anatoly C2011, showed outstanding high yielding in most environments and were stable depending on at least one methodology. These commercial cultivars should be prioritized for promotion to farmers to support Mexican wheat productivity.

## ACKNOWLEDGEMENTS

The authors acknowledge the support given by Fondo Sectorial SAGARPA-CONACYT through Project No. 146788 “Sistema de mejoramiento genetico para generar variedades resistentes a royas, de alto rendimiento y alta calidad para una produccion sustentable de trigo en Mexico”, CIMMYT and the Secretaria de Agricultura y Desarrollo Rural (SADER) through the MasAgro Initiative and CONACYT for providing a postgraduate scholarship (no. 374086) to J. L. V. A. F. J. P.-C. and A.R.B. are supported by the One CGIAR Accelerated Breeding Initiative.

## SUPPLEMENTARY MATERIAL

**Supplementary Table S1.** Combined variance components, broad-sense heritability, overall gran mean for grain yield (t ha^-1^) of bread and durum wheat in trials in 2012/13 and 2013/14 in northwest, north and central Mexico regions.

| Source of variation | Variance |
| --- | --- |
| Environment (E) | 2.377*** |
| Genotype (G) | 0.052*** |
| G × E | 0.201*** |
| Error | 0.38 |
| Heritability | 0.79 |
| Grand mean | 4.96 |
| Coefficient of variation (%) | 12.4 |

\*\*\*, Statistically significant with $P < 0.001$ .

**Supplementary Table S2.** Summary of the analysis of variance of the Eberhart and Russell (1966) stability parameters

| Source of variation | Degrees of freedom | Sum of Squares | Mean square |
| --- | --- | --- | --- |
| Genotype (G) | 15 | 13.80 | 0.92*** |
| Environment (linear) | 448 | 1248.27 | 2.79 |
| Environment + G × E | 1 | 1084.00 | 1084 |
| G × E (linear) | 15 | 87.46 | 5.83*** |
| Pooled deviation | 432 | 76.81 | 0.18 |
| **Bread wheat** |  |  |  |
| Bacorehuis F2015 | 27 | 3.06 | 0.11 |
| Borlaug100 F2014 | 27 | 3.65 | 0.14 |
| Conatrigo F2015 | 27 | 1.74 | 0.06 |
| Villa Juarez F2009 | 27 | 8.54 | 0.32* |
| Kronstad F2004 | 27 | 5.24 | 0.19 |
| Roelfs F2007 | 27 | 3.24 | 0.12 |
| Tacupeto F2001 | 27 | 6.36 | 0.24 |
| Urbina S2007 | 27 | 7.93 | 0.29* |
| **Durum wheat** |  |  |  |
| Barobampo C2015 | 27 | 4.15 | 0.15 |
| CONASIST C2015 | 27 | 4.32 | 0.16 |
| Anatoly C2011 | 27 | 4.80 | 0.18 |
| CIRNO C2008 | 27 | 3.78 | 0.14 |
| Gema C2004 | 27 | 2.47 | 0.09 |
| Cevy Oro C2008 | 27 | 7.09 | 0.26 |
| Sawali Oro C2008 | 27 | 5.73 | 0.21 |
| Movas C2009 | 27 | 4.71 | 0.17 |
| Pooled error | 464 | 82.68 | 0.18 |

\*,\*\*\* Statistically significant with $P < 0.05$ and $< 0.001$ .

